# Optimized AAV capsids for diseases of the basal ganglia show robust potency and distribution in adult nonhuman primates

**DOI:** 10.1101/2024.05.02.592211

**Authors:** DE Leib, YH Chen, L Tecedor, PT Ranum, MS Keiser, BC Lewandowski, EM Carrell, S Arora, I Huerta-Ocampo, X Liu, BL Davidson

## Abstract

Huntington’s disease and other disorders of the basal ganglia create challenges for biomolecule-based medicines given the poor accessibility of these deep brain structures following intracerebral or intravascular delivery. Additionally, for adeno-associated viruses (AAVs) intravascular delivery exposes peripheral tissues to the vast majority of the therapy, increasing the risk of immune responses and the quantity and associated cost of goods required for therapeutically relevant brain penetration levels. Here, we found that low dose, low volume delivery of unbiased AAV libraries into a focused brain region allowed recovery of novel capsids capable of broad access to key deep brain and cortical structures relevant for human therapies at doses orders of magnitude lower than used in current clinical trials. One such capsid, AAV-DB-3, provided transduction of up to 45% of medium spiny neurons in the adult NHP striatum, along with substantial transduction of relevant deep layer neurons in the cortex. Notably, AAV-DB-3 behaved similarly in mice as in NHPs and also potently transduced human neurons derived from induced pluripotent stem cells. Thus, AAV-DB-3 provides a unique AAV for network level brain gene therapies that translates up and down the evolutionary scale for preclinical studies and eventual clinical use.

## Introduction

Recombinant adeno-associated viruses (rAAVs) are a promising therapeutic modality to treat neurodegenerative diseases for their high safety profile, ability to infect non-dividing cells and to facilitate long-term transgene expression. Naturally occurring serotypes are effective at targeting different cells of the nervous system with variable tropism determined by capsid identity and route of administration^1, 2^. Broad transduction of the central nervous system (CNS) can be achieved through non-invasive intravenous (IV) delivery of AAV9 and AAVrh10 and numerous modified capsid variants designed to cross the blood-brain barrier (BBB)^3-9^. However, this approach lacks specificity and has low targeting efficiency with substantial peripheral virus burden that can contribute to immune responses and liver toxicity^10-14^. Direct, intraparenchymal delivery of AAV confers higher specificity and targeting efficiency, but tissue distribution is largely limited to the site of injection with large brain structures requiring multiple injections^15-17^ and methods to enhance delivery^2, 18^.

Huntington’s disease^19^ is an autosomal dominant disorder characterized by progressive degeneration and transcriptional dysregulation in the basal ganglia with subsequent changes in cortical regions of the brain^20-22^. The basal ganglia comprise a collection of ganglia and nuclei located deep within the brain that form connections important for movement, decision making, and reward behaviors. Major functional components include the caudate nucleus, globus pallidus, putamen, substantia nigra, subthalamic nucleus and ventral pallidum. The globus pallidus (“pale globe”) represents a central figure in the basal ganglia circuitry and is divided into two functionally distinct segments, internal (GPi) and external (GPe). GPe receives input from the striatum (caudate and putamen) and projects outwards towards the subthalamic nucleus. GPi also receives input from the striatum and projects to the thalamus, a projection that enables indirect connection with cortical regions of the brain. Together, affected regions in HD pose unique challenges for therapeutic delivery: relevant structures lie deep within the brain and span large areas that cannot be fully targeted by a single, direct delivery.

Despite these challenges, much effort has been put toward development of therapies for HD, including strategies to knockdown mutant huntingtin (Htt) gene product or indirectly impact disease pathogenesis^23-27^. Preclinical studies in mice have shown therapeutic benefit following a single, direct injection of AAV into the striatum, however this paradigm is not suited for the larger structures of the NHP or human brain with currently available capsids. An ongoing clinical trial to treat HD targets huntingtin transcripts (*HTT*) with an AAV5-delivered artificial miRNA at three sites in the caudate nucleus and putamen per hemisphere using convection-enhanced delivery^28^ (NCT04120493).

To overcome the challenges for AAV gene therapies for disorders that impact cortical and subcortical structures like HD^21, 29^, we devised an unbiased approach to identify highly potent capsids capable of transducing our target brain regions following delivery to the globus pallidus. Our libraries were based on varying AAV serotypes, required only several animals for lead selection, and the resulting capsids did not require further optimization to improve their trafficking and targeting capabilities. Notably, our most potent capsid, AAV-DB-3, was greater than two orders of magnitude more potent than capsids being assessed clinically for HD. AAV-DB-3 also efficiently transduces human iPSC-derived neurons and retains the properties of broad distribution across mice strains, supporting their application to pre-clinical testing and relevant dose range finding studies important for advancing to HD patients.

## Results

### *In vivo* screen of a peptide modified AAV library

A peptide modified AAV capsid library approach was leveraged to identify highly performant capsids variants targeting deep brain structures after low dose infusion into the globus pallidus (GP). Three parental serotypes (AAV1, AAV2, and AAV9) were modified to create the capsid variant libraries used in this work. Each parental serotype was modified by insertion of a 7-mer amino acid sequence into the variable region of loop 8 at positions 590, 587, and 588 in AAV1, AAV2, and AAV9 respectively. The insertions are a sequence of semi-random DNA bases “NNKNNKNNKNNKNNKNNKNNK” where “N” represents DNA bases A, C, T or G and “K” represents G or T. Our AAV selection methodology consisted of several rounds of *in vivo* selection in adult NHPs accompanied by Illumina amplicon sequencing to resolve and quantify enrichment of capsid variants (Figure 1A). The estimated diversity of the initial screening library used in this work was ∼6.8 million. After two rounds of *in vivo* target tissue enrichment screening, the total number of remaining capsid variants across brain regions of interest was 42,487, a reduction of more than two orders of magnitude from the initial screening library. To assess individual capsid variant performance, amplicon sequencing was performed resulting in the identification of 161 top-performing capsid variants based on broad expression across various brain regions and/or selective enrichment in target regions (caudate nucleus and putamen) (Figure 1B, C). A final round of screening using this small library of top performing capsids was performed in both an adult rhesus macaque and African green monkey to assess cross-species performance. From analysis of amplicon sequencing data of this final screen, we selected 6 capsids (2 variants for each serotype) for further validation based on performance across brain regions relevant to HD.

**Figure 1.**
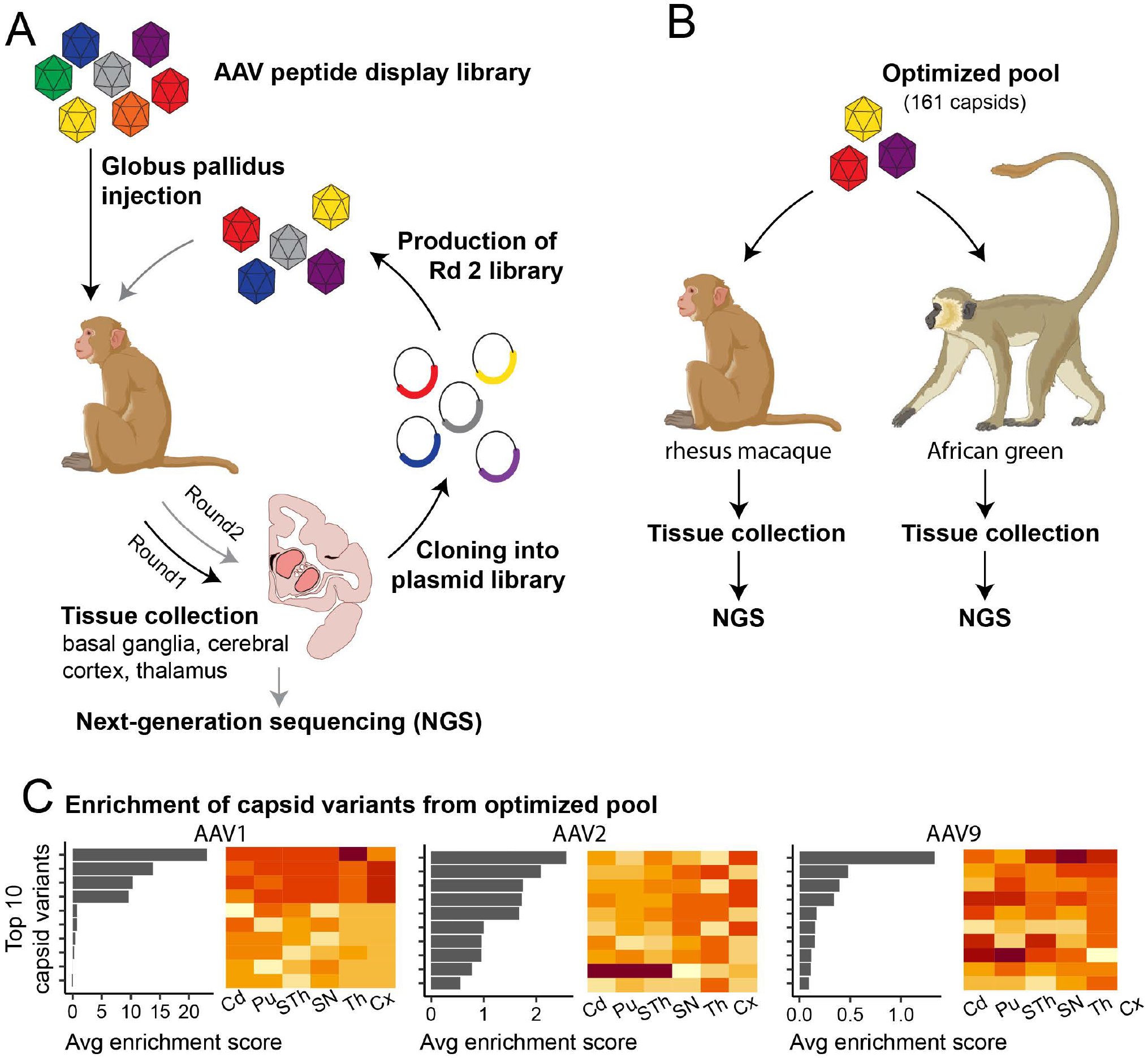
AAV capsid library enrichment and candidate selection A) Schematic depicting the AAV capsid selection pipeline used in this work. AAV peptide display libraries were designed to contain a semi-random heptapeptide insert in multiple parental AAV serotypes (AAV1, AAV2, and AAV9), yielding an initial ∼6.8 million capsid variants. Enrichment and selection of capsids is achieved by sequential passages through primate brain. The first round of enrichment (black arrows) involved a single infusion of ‘Round 1’ input library composed of peptide-modified AAV capsids from three parental serotypes. The input library was injected by bilateral intraparenchymal (IP) infusion into the globus pallidus of a rhesus macaque. After two days in life, CNS regions of interest were collected. DNA was extracted from collected tissues and recovered AAV variant sequences were used to generate a new (Round 2) AAV input library. A second round of screening (grey arrows) was initiated by bilateral GP infusion of the Round 2 input library into rhesus macaque. For round 2 screening, both viral DNA and RNA were collected from tissues. Peptide modifications of successfully enriched capsids were identified using Illumina sequencing of viral amplicons. Capsid performance was assessed using a combination of total detection in target tissues and ‘normalized enrichment’: the ratio of a capsid’s relative abundance in a target region vs the input library. B) 161 top ranking capsids across all parental serotypes were identified to generate a third input vector library hereafter referred to as the “optimized pool”. A final round of screening was initiated by injection of the optimized pool input vector into the GP an adult rhesus macaque and African green monkey. Tissues collected from these primates became the basis for DNA- and RNA-based amplicon sequencing. C) Six highly enriched capsid variants were identified based on a combination of overall detection across brain regions, and/or targeted enrichment in caudate and putamen, and were selected for fluorescence validation. Selected capsids expressing fluorescent payloads were individually cloned and produced. They were then pooled together for subsequent infusion into additional primates for small pool fluorescent validation.

### AAV-DB candidate validation in adult NHPs

The deep-brain targeting capsid variants (AAV-DB) selected for further validation were packaged with distinct reporter transgenes driven by the ubiquitous CAG promoter. Three AAV-DB candidates expressing different fluorescent payloads were mixed and injected at a dose of 7.5E10 vg each into the GP of an adult rhesus macaque (Fig. 2A). After approximately 3 weeks the animal was euthanized, and tissue processed for histology. Expression of fluorescent reporters was assessed using direct fluorescence *in situ* hybridization (FISH) for their respective transcripts (Fig. 2B). AAV -DB-1, 2 and 3 transduced numerous cells in the GP, putamen, and caudate, with particularly robust transduction by AAV-DB-2 and DB-3 (Fig. 2C). For AAV-DB-2 and DB-3, transduction was remarkably widespread in the caudate nucleus and putamen, reaching distances greater than a centimeter from the infusion site. AAV-DB-1, DB-2, and DB-3 also transduced cells in the substantia nigra reticulata (Fig. 2C vii,viii).

**Figure 2.**
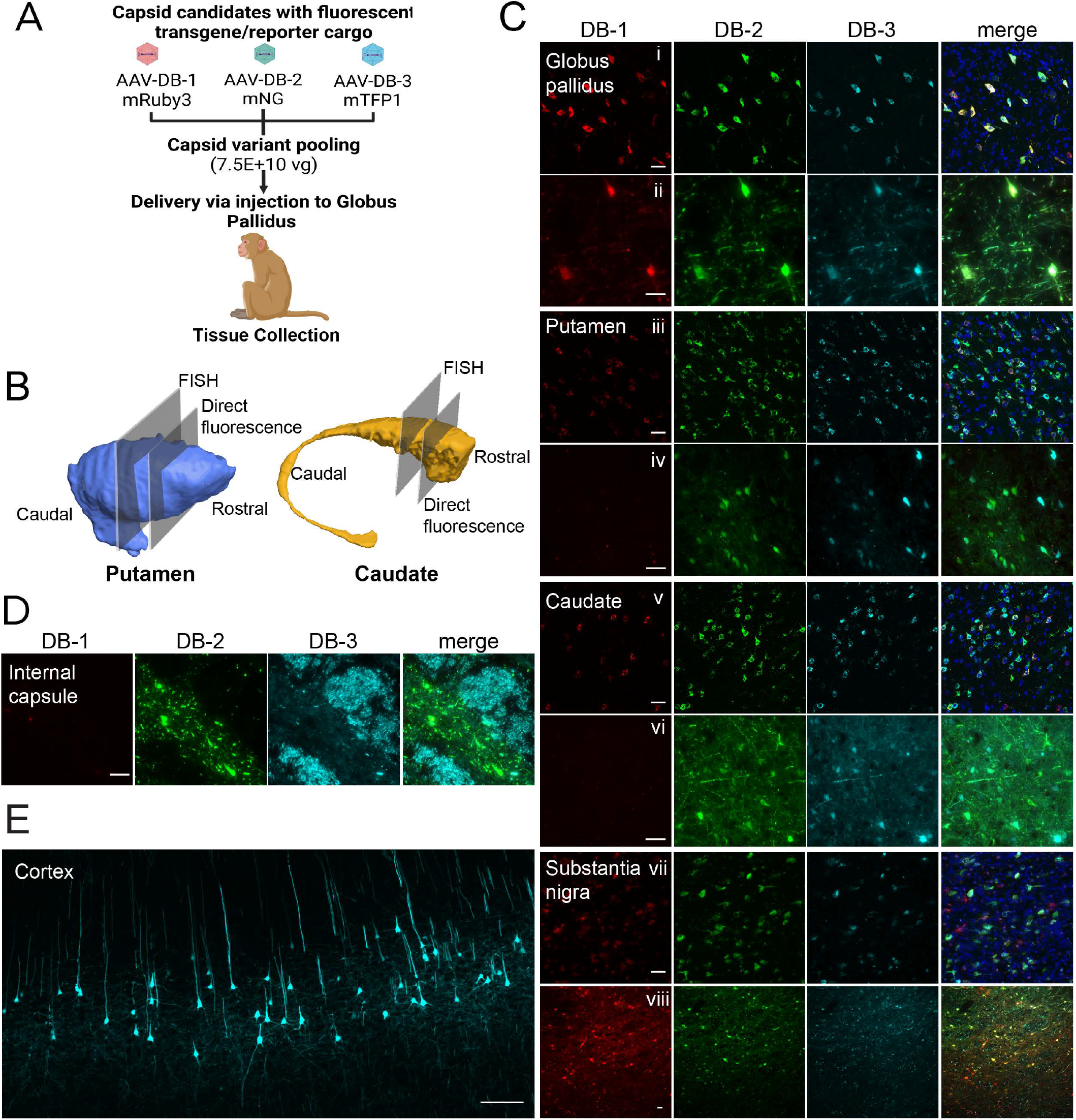
AAV variant validation. A) A mix of DB-1.mRuby3, DB-2.mNG, and DB-3.mTFP1 AAV variants was infused into the globus pallidus of a rhesus macaque at 7.5E10 vg/capsid. B) Renderings of the NHP putamen and caudate^43^ are intersected with planes indicating the anatomical positions for direct fluorescence or FISH analysis. C) AAV-DB-1, AAV-DB-2, and AAV-DB-3 transduction in the globus pallidus (i, ii), putamen (iii, iv), and caudate (v, vi). Cell transduction was analyzed by FISH (i, iii, v, vii), and direct fluorescence of expressed fluorophores (ii, iv, vi, viii). D) mTFP1 fluorescence in the fibers of the internal capsule. E) Direct fluorescence of mTFP1 positive neurons in motor and premotor cortex. Scale bar: A-C 50 µm, D 200 µm.

In tissues outside the basal ganglia, AAV-DB-3.mTFP1 robustly transduced layer 5/6 subcortical projection neurons in cerebral cortices (Fig. 2E). Subcortical projection neurons are particularly vulnerable in HD, and transduction of these cells therefore makes AAV-DB-3 a particularly attractive capsid variant for HD gene therapies^21^. As further confirmation of cortical transduction, we also observed abundant AAV-DB-3.mTFP1 signal in axons within the internal capsule (Fig 2D). There were scattered transduced cells in other brain areas, including thalamus and the cingulate cortex (Fig. 3A), but transgene expression was largely confined to the basal ganglia and connected brain areas and limited predominately to the ipsilateral side of the infusion (Fig. 3B). AAV-DB candidates were also mixed and injected at a lower dose of 3.0E10 vg each into another adult rhesus macaque. AAV-DB-3 performed similarly at this lower dose, showing widespread transduction in the striatum and motor cortex (Fig. 3C). There was little to no transduction observed in dorsal root ganglia or liver, which are common off-target tissues for CSF-or IV-delivered AAVs^10, 30, 31^ (Suppl. Fig. 1). Based on AAV-DB-3.mTFP1’s widespread transgene expression across basal ganglia regions and cerebral cortex and its limited to undetectable off-target tissue transduction, we selected AAV-DB-3 as the top AAV capsid variant for deep brain applications and further testing in mice strains and neurons derived from human iPSCs.

**Figure 3.**
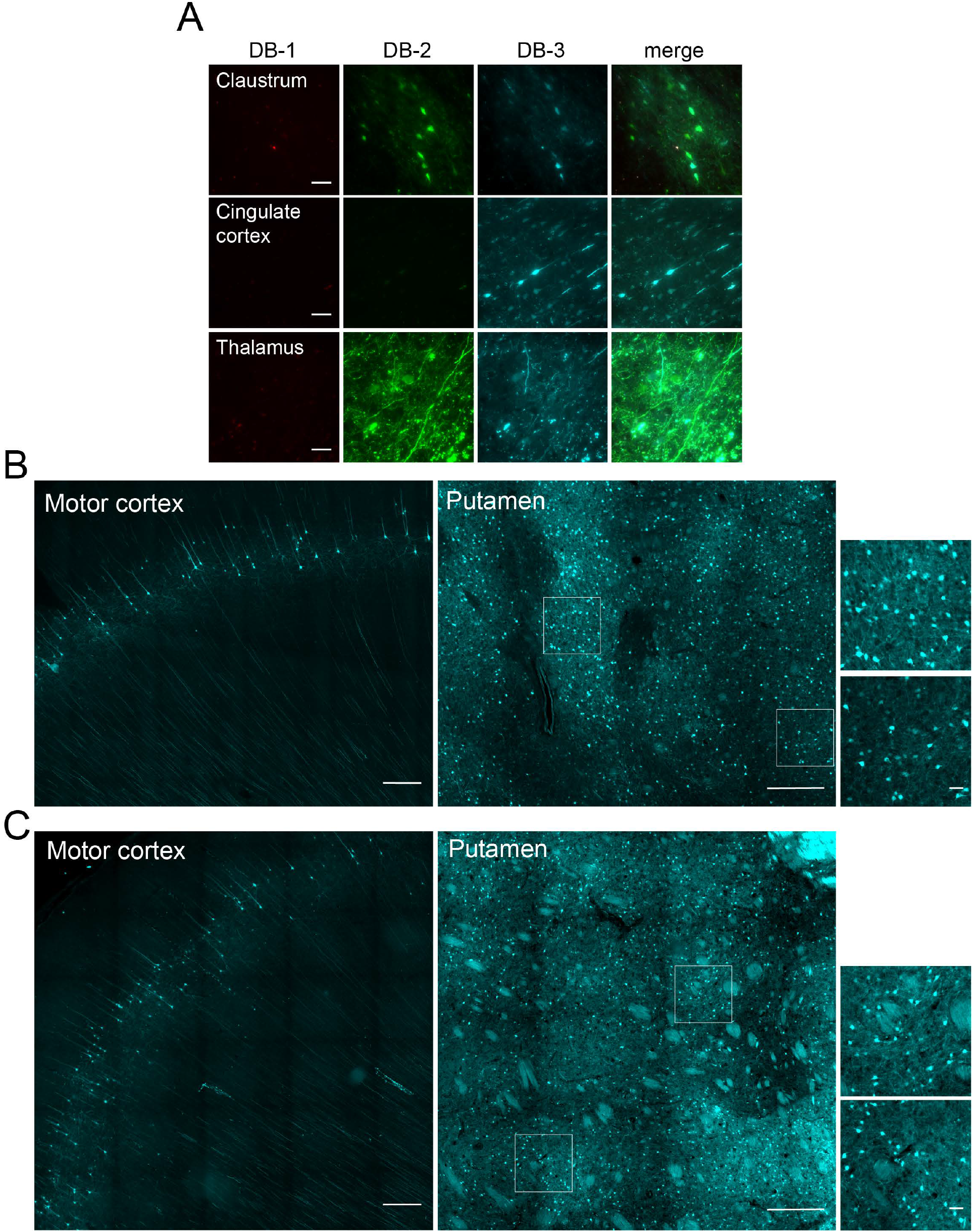
AAV-DB-1, DB-2, and DB-3 biodistribution in CNS after GP injection. A) Distribution of AAV-DB-1, AAV-DB-2, and AAV-DB-3 in the claustrum, cingulate cortex, and thalamus marked by direct fluorescence of packaged fluorophores. B,C) Tile scans showing the extent AAV-DB-3.mTFP1 biodistribution in motor cortex and putamen following injection of 7.5E10 (B) or 3E10 (C) vg into the GP. Scale bar: A) 50 µm, B,C) 500 µm

To further examine cell types transduced by AAV-DB-3.mTFP1, we performed co-labeling experiments by RNA-FISH focusing on neurons in the caudate nucleus and putamen, as medium spiny neurons (MSNs) are among the more vulnerable neurons in HD and a critical cell type for other basal ganglia diseases. RNA FISH for mTFP1 and co-labeling with the neuronal marker gene NeuN revealed that the majority of the transduced cells in the caudate and putamen were neurons (Fig. 4A). Two major classes of MSNs, D1-MSNs and D2-MSNs, express D1 or D2 dopamine receptors (encoded by *DRD1* and *DRD2*, respectively). RNA FISH for mTFP1, *DRD1*, and *DRD2* showed that dopamine receptor expressing MSNs in the caudate and putamen were the most highly transduced cell type; 55.6% of D1-MSNs and 31.6% of D2-MSNs co-expressed mTFP1, reaching a combined of 40.4% of all MSNs transduced (Fig. 4B, C). Note that the doses used in these animals were relatively low (3E10 or 7.5E10 total vg) compared to conventional viruses like AAV2 or AAV5 applied using convection enhanced delivery^18, 32^, which was not required here given the low volumes of infusate. Higher doses may provide even greater coverage of MSNs. Taken together, transgene expression data in adult, old-world NHPs showed potent transduction of a significant percentage of neurons of the basal ganglia and cerebral cortex after a single, low-dose infusion of AAV-DB-3.

**Figure 4.**
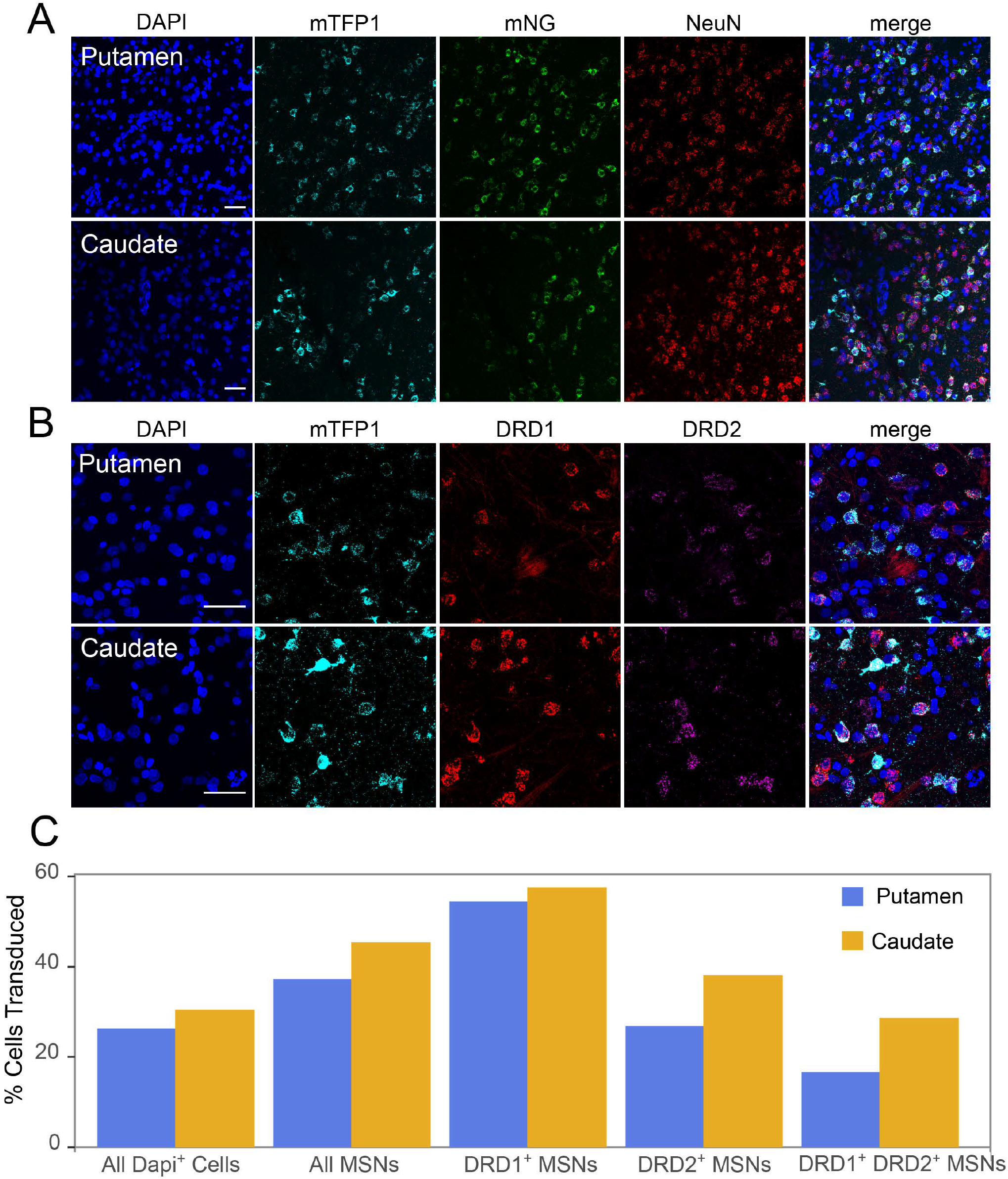
Characterization of AAV-DB-2 and AAV-DB-3 transduced cells in NHP caudate and putamen. A) FISH detection of the neuronal specific marker NeuN and mTFP1 or mNG transcripts in DB-2 or DB-3 transduced cells, respectively, in putamen and caudate. B) FISH analyses for DRD1 or DRD2 medium spiny neuron (MSN) subpopulations also transduced by AAV-DB-3.mTFP1. C) Quantification of the percentage of transduced cells among the indicated MSN subtypes. MSNs counted from images taken across the ventral-dorsal axis of the putamen and caudate (Figure 2B).

### AAV-DB-3 transduction properties in mice

To evaluate the evolutionary conservation of AAV-DB-3, we injected C57/BL6 mice in the mouse equivalent of the external globus pallidus (GPe) with AAV-DB-3.mTFP1. Three weeks after injection, brain tissue was harvested for histological analysis. In addition to abundant GPe neurons transduced around the injection site (Fig. 5A, B), we observed mTFP1 expression in numerous MSNs in the striatum (Fig. 5C). We also observed abundant neuronal transduction in the motor, cingulate and medial prefrontal cortices (Fig 5A, D). These results largely mirror the NHP transgene expression results and support an evolutionarily conserved mechanism for AAV-DB-3 biodistribution. In parallel, two groups of C57/BL6 mice were injected in the striatum or thalamus (Suppl. Fig 2). The brains were collected, and tissue sections were processed in the same manner as above. Our results after striatal injections show a vast number of labeled striatal MSNs (Suppl. Fig. 2B), pallidostriatal neurons in the GPe (Supp. Fig. 2F) and numerous neurons located in somatosensory, frontal and motor cortices (Suppl. Fig. 2E), as well as in thalamostriatal neurons (Suppl. Fig. 2C, D). In mice injected in the thalamus (Supp. Fig. 2G), we observed numerous neurons located in several thalamic nuclei (i.e, reticular nucleus, geniculate thalamus, and the ventral thalamus. Suppl. Fig. 2H, J), as well as in the somatosensory, motor and frontal association cortices (Suppl. Fig. 2I) and the substantia nigra (Suppl. Fig. 2K).

**Figure 5.**
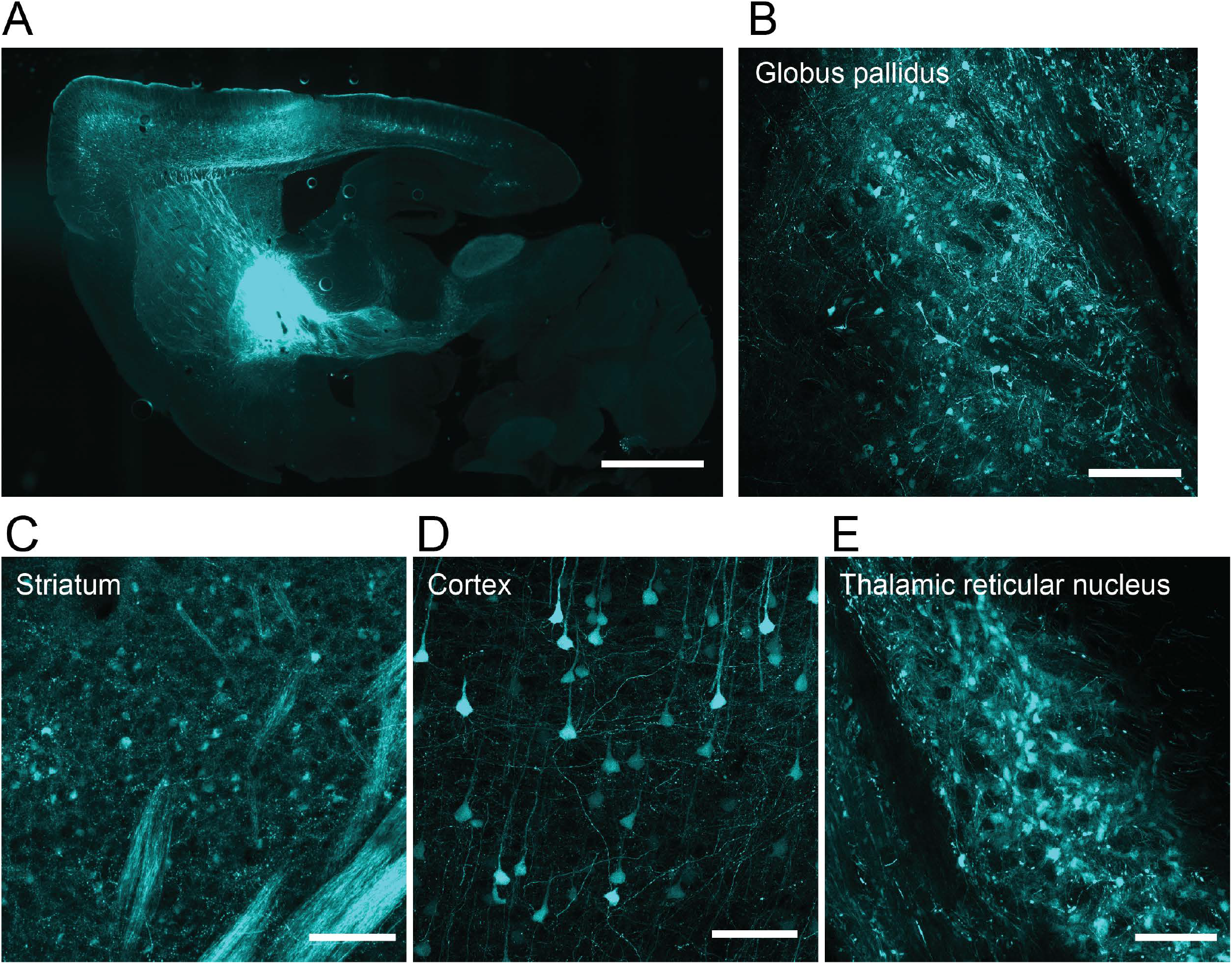
Distribution of AAV-DB3-mTFP1 in mouse basal ganglia and connected structures. A) Schematic of the procedure. B, mTFP1-positive cells in the GPe. C, Retrogradely transduced mTFP1^+^ MSNs in the striatum. D, Transgene positive neurons in the somatosensory and motor cortices. E, mTFP1 positive neurons in the reticular thalamic nucleus. Scale bar: A) 1.5mm. B-E) 200 microns.

### AAV-DB-3 efficiently transduces human neurons

We further evaluated the transduction efficiency of AAV-DB-3 in human neurons derived from induced pluripotent stem cells. iPSC-derived cortical neurons were transduced at day 40 of differentiation with AAV-DB-3.mTFP1 or its parental serotype AAV1-eGFP at multiple multiplicities of infection (MOIs; 1E3, 1E4 and 1E5). On day 10, qualitative analysis by microscopy revealed significantly higher transduction by AAV-DB-3 compared to AAV1 at equivalent doses (Fig. 6A). Moreover, at the lowest MOI of 1E3, AAV-DB-3 showed a 20-fold increase in transgene expression by RT-qPCR compared to the wild-type parental serotype (Fig. 6B). AAV-DB-3 may therefore be a valuable tool to manipulate gene expression in cultured human neurons and further supports its use in pre-clinical testing for human disease application.

**Figure 6.**
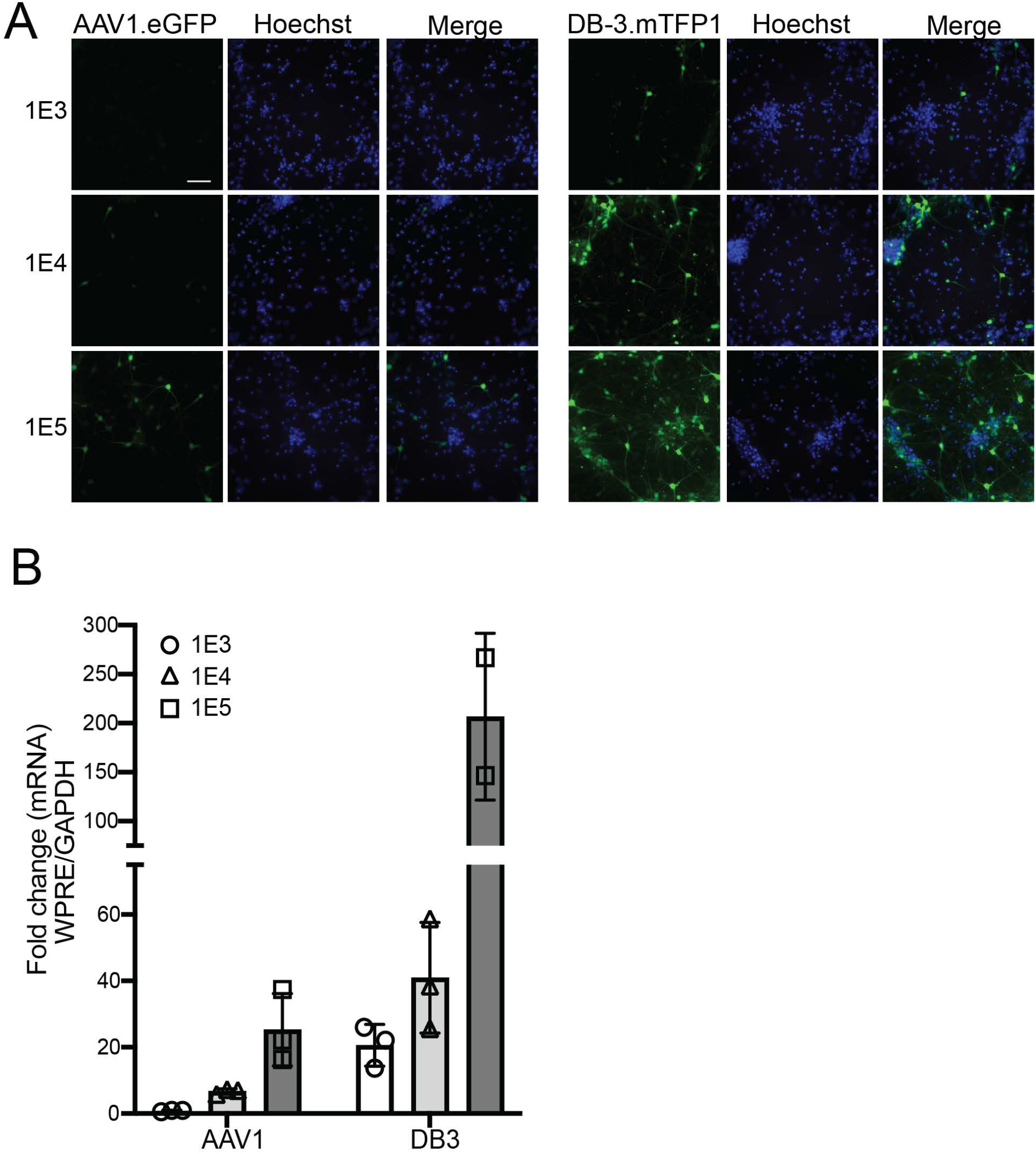
Transduction of human iPSC derived neurons by AAV1.eGFP and AAV-DB-3.mTFP1. A) Direct fluorescence of AAV1.eGFP and AAV-DB-3.mTFP1 in iPSC-derived excitatory neurons at 10 days post-transduction with indicated MOIs. Nuclei were stained with Hoechst 3325. Scale bar is 50 µm and applies to all panels. B) Quantitative PCR for transgene expression. All samples were normalized to *GAPDH* and are presented relative to AAV1.eGFP at the 1E3 dose.

## Discussion

AAV gene therapies for disorders with dysfunction in deep brain areas have shown early promise despite the challenges with delivery to these regions. Of note, in children with aromatic L-amino acid decarboxylase (AADC) deficiency, delivery of an AAV2 vector expressing AADC to the substantia nigra and ventral tegmental area was well tolerated and led to clinical improvements^33^. In PD patients, AAV2 gene therapies with AADC or glial cell derived neurotrophic factor (GDNF) payloads delivered to the putamen were well tolerated and, for AADC, showed possible stabilization of motor function^17, 34^. A gene therapy for HD with an AAV5 vector expressing a huntingtin-lowering microRNA delivered to the caudate and putamen is also in clinical trials (NCT04120493). However, these approaches rely on large infusion volumes and convection enhanced delivery via multiple cannula trajectories per hemisphere to achieve adequate deep brain coverage. For example, in the AAV2 AADC trial, patients in the highest dose group received 900 µL of test article per hemisphere via 2-3 cannula trajectories, and coverage of the putamen was around 42% based on intraoperative MRI scans^17^. These injection volumes necessitate long surgical times, typically under MRI guidance, increasing patient burden and creating logistical challenges when scaling a therapy to a larger number of patients.

An alternative to direct intraparenchymal delivery of gene therapies to deep brain areas is IV-delivery of AAVs with engineered capsids that cross the blood-brain barrier (BBB). Naturally occurring AAV variants poorly penetrate the BBB, with most viral particles accumulating in the liver and other peripheral organs^14, 30, 35^. In mice, peptide-modified AAV9 capsid variants were identified with a dramatically enhanced ability to cross the BBB^5, 6^. However, this ability was not retained in all mouse strains or in nonhuman primates^36-38^. Building on this work, new AAV variants with greater potential for clinical application have been identified through a variety of approaches, including *in vivo* screens in NHPs and human transferrin receptor binding^8, 9, 39^. These efforts show promise, and progress continues towards identifying more potent BBB-penetrant AAV capsids. If sufficiently potent capsids are developed, challenges remain for IV delivery including safety concerns around peripheral exposure to high doses of AAV^12, 13^ as well as manufacturing the high doses required.

Here, we identified capsid variants capable of broad transduction of neurons in deep brain areas in NHPs following low volume intraparenchymal infusion in the GP. The most potent variant, AAV-DB-3, transduced cells across multiple basal ganglia and cortical regions in NHPs, including MSNs in the caudate nucleus and putamen. AAV-DB-3 also potently transduced neurons in the mouse brain as well as human IPSC-derived neurons in culture. The cross-species validation of AAV-DB-3 in two NHP species, mice, and human neurons contrasts with results for other engineered capsids that show highly species-specific performance^9, 36^. The pattern of AAV-DB-3’s spread in the NHP and mouse brain point to retrograde transport along axons, but additional studies will be required to characterize the mechanism of AAV-DB-3’s remarkable spread in the brain. Of note, AAV-DB-3’s parental serotype is AAV1 and not AAV9, commonly used as the sole serotype for similar capsid screens, and AAV-DB-3 would not have emerged if our initial capsid screen did not contain multiple serotypes in addition to the commonly used AAV9. Taken together, our results suggest that delivery of AAV-DB-3 to the GP may provide an alternative to high-volume convection enhanced delivery or high-dose IV delivery for AAV gene therapies targeting deep brain areas. Infusions in the GP are expected to be clinically feasible, as functional neurosurgeons have been placing electrodes in the GPi for deep brain stimulation for 30 years^40^. One potential application for AAV-DB-3 is HD, for which various cargos may be packaged targeting *HTT* or other implicated genes including *MSH3* or *PIAS1*^*26*, *27*, *41*^.

## Acknowledgements

We thank Laurence Busque, Ashley Robbins, Melanie McFadden and the members of the CHOP Comparative Medicine Services Core and ClearPoint team members for their contributions to this work.

## Methods

### Peptide modified AAV library description and construction

Plasmids used for the AAV library construction were generated by cloning AAV2 Rep and AAV Cap of different serotypes into an ITR-containing plasmid using Gibson assembly. Two unique restriction enzyme (RE) sites were built flanking the peptide insertion site in the AAV Cap gene. Oligonucleotides encoding the random 7-mer (NNK)_7_ were synthesized by Integrated DNA Technologies (Coralville, IA). The oligonucleotide library was cloned into the RE digested recipient plasmids using Gibson assembly. Pooled plasmids were transformed into Endura electrocompetent cells (LGC, Middlesex, UK) and plasmid isolation performed using Qiagen Giga Prep Endofree kits (QIAGEN, Germantown, MA). For the second round library generation, viral genomic DNA from collected tissues was isolated by QIAamp DNA mini kits (QIAGEN, Germantown, MA) and the variable regions of the AAV library amplified using serotype-specific primers and the NEB Q5 2x Master Mix (New England Biolabs, Ipswich, MA). Amplicons were incorporated into recipient plasmids by Gibson assembly and transformed into Endura electrocompetent cells (LGC, Middlesex, UK).

### AAV library virus production

AAVs were generated in HEK 293 cells transfected with pAd helper and pAAV.Lib plasmids, the latter maintained at 1000 plasmid copies per cell. Recombinant virus was harvested and purified by two rounds of cesium chloride gradient centrifugation and AAVs titered by ddPCR.

### NHP animals and procedures

This study was conducted in accordance with the Guide for the Care and Use of Laboratory Animals (National Research Council) and all procedures were approved by the CHOP Research Institute Animal Care and Use Committee. Animals were housed at the CHOP Research Institute under a 12-hour light:dark cycle with ad libitum access to purified drinking water and twice daily feedings with Purina LabDiet Certified Primate Diet (5048) enriched with fruits and vegetables. On the day of surgery, the GP was targeted by ClearPoint MRI-guided infusion using a 3T research MRI unit (Siemens Trio; SYNGO Vb17). A ClearPoint SmartFlow cannula was attached to a BD syringe controlled by a Harvard Apparatus infusion pump. The cannula was primed with test article and inserted into the brain to the desired depth before infusing 30 uL of test article at a rate of 1.0 uL/min. Following the infusion and a 10-minute dwell period, the cannula was removed, the skin sutured, and the animal recovered.

AAV9.eGFP (3E13 vg total), a mid-range dose for CSF-delivered AAV9^31^ was infused as a positive control for DRG transduction^10^. Infusion was to the left lateral ventricle using a stereotactic device for NHPs using coordinates calculated from baseline MRI scans and confirmed by administration of a small volume of Iso-Vue M contrast under fluoroscopic guidance. After a three-minute dwell period, the needle was removed, the skin sutured, and the animal recovered. Buprenorphine SR was given as analgesia, and the animals were monitored postoperatively for pain and welfare.

NHPs were euthanized by exsanguination. Animals were sedated, transcardially perfused with ice-cold saline, and the brain removed and processed into 4-mm coronal slabs in a rhesus macaque brain matrix. For AAV library experiments, tissue samples from basal ganglia, thalamus, cerebral cortex, and other brain areas were micro-dissected and flash frozen for analysis. For fluorophore experiments, brain slabs, DRGs, and other tissues were postfixed in 4% paraformaldehyde. Additional samples for molecular analysis were banked, including liver samples used for ddPCR.

### Amplicon sequencing of AAV libraries

To quantify input library composition and viral transduction in target tissues, amplicons were created from the peptide modified region of the viral Cap gene and sequenced. Genomic DNA (gDNA) or RNA from the tissues was isolated using a QIAamp DNA mini kit or RNeasy Plus kit (QIAGEN, Germantown, MA). Sequence encoding the modified capsid insertion was amplified using serotype-specific primers with PCR cycles less than 30 by using Q5 2x Master Mix (NEB, Ipswich, MA). Two microliters of 1^st^ round of PCR products were input to total 50 µl reaction of second round PCR to introduce Illumina i5 and i7 sequencing indices. PCR products were run ona 3% agarose gel to recover of the NGS amplicon libraries and purified by MinElute Gel Extraction Kit (QIAGEN, Germantown, MA). Libraries were sequenced on an Illumina Novaseq 6000 using a 350-cycle reagent kit and paired end read chemistry yielding 175 bp paired end read outputs. Target read depth was variable and allocated according to library identity and diversity. Read depth targets ranged from 2 to 10 million reads per sample.

### ddPCR for AAV genomes

DNA was isolated from NHP liver samples using the QIAamp DNA kit (Qiagen). ddPCR was performed on the isolated DNA using the ddPCR Supermix for Probes (BioRad) on the QX200 ddPCR instrument (BioRad) with primer-probe sets for the CAG promoter in each fluorophore transgene and NHP gDNA. Values were averaged from two technical repeats.

### Mouse biodistribution studies

All procedures were approved by the Children’s Hospital of Philadelphia Institutional Animal Care and Use Committee. Mice were maintained on a 12 h light/dark cycle and ad libitum access to water and food. Adult C57BL/6 mice (n=3) were injected with AAV-DB-3.mTFP1 at 1E10 vg. Injections targeting the GPe (from bregma in mm: AP, -0.46; ML, ±1.85; DV, 3.7; n=3) were done at a rate of 25 nl/min. Injections targeting the thalamus (from bregma in mm: AP, -1.70; ML ±1.25; DV, 3.3; n=3) or striatum (from bregma in mm: AP, +0.5; ML ±1.75; DV, 3.2, 2.6; n=3) were performed using the same dose and injection rate. Three weeks after injection, mice were transcardially perfused with 0.05 M PBS, pH 7.4, followed by 50 ml of 4% w/v paraformaldehyde in phosphate buffer (0.1 M, pH 7.4). Brains were cryoprotected and sagittally sectioned. For each brain, the sites of injection were verified, and sections analyzed using a Leica DM6000B epifluorescence microscope for low power and tiled images. Z-stacks were acquired using a Leica SP8 confocal microscope.

### RNA-FISH

4 mm coronal NHP brain slabs were post-fixed in 4% PFA, sucrose cryoprotected, and OCT embedded before 16 µm thick slices were collected on Superfrost Plus slides. Fluorescent *in situ* hybridization was performed using RNAscope Multiplex Fluorescent Reagent Kit v2 Assay (Advanced Cell Diagnostics, Cat. 323100-USM) following manufacturer guidelines.

### RNA-FISH quantification of MSN transduction

Three RNAscope probes recognizing DRD1 (Advanced Cell Diagnostics Cat. 549041), DRD2 (Cat. 549031-C2), and mTFP1 (Cat. 500271-C3) were used to label regions of the caudate and putamen from two NHPs. Specific Opal fluorophores were associated to each RNAscope probe (DRD1-C1-Opal520, DRD2-C2-Opal570, and mTFP1-C3-Opal620). Forty-two total images were collected from these regions and quantified using an unbiased automated cell counting approach. Puncta in all channels greater than .002 µm^2^ were detected using QuPath with the following relative fluorescence units (RFU) minimum pixel intensity thresholds DRD1 (Opal520) ≥25, DRD2 (Opal570) ≥25, mTFP1 (Opal620) ≥ 50. Detected puncta were then counted and counts tables were processed using a custom R computational pipeline. Nuclei were defined as being DRD1 or DRD2 positive neurons if they contained ≥ 50 puncta in the respective channel. If nuclei contained >60% puncta of either Opal520 or Opal570-positive they were assigned that DRD1 or DRD2 identity accordingly. If a nucleus containing >= 50 puncta did not meet this threshold it was classified as DRD1 and DRD2 double positive. Nuclei of all types were defined as positive for mTFP1 transduction if they were found to contain >=2 mTFP1 puncta. The above settings were selected to achieve consistency with human counting results and for their ability to clearly differentiate positive nuclei from non-cellular background signal.

### Human iPSC-derived neuron transduction

iPSC-derived cortical excitatory neurons were generated from iPSCs as previously described^42^. Virus transduction was performed at Day 40 of differentiation during a scheduled half-media change. Ten days later RNA was isolated for qRT-PCR using standard methods. For microscopy, iPSC-derived cortical neurons were seeded on coverslips, and transduced as above and 10 days later cells were fixed, blocked, and fluorescence levels assessed using epi-fluorescent microscopy (Leica, DM6000B).

