## Supplemental Figures for "Optimized AAV capsids for diseases of the basal ganglia show robust potency and distribution in adult nonhuman primates"

Supplemental Figure 1

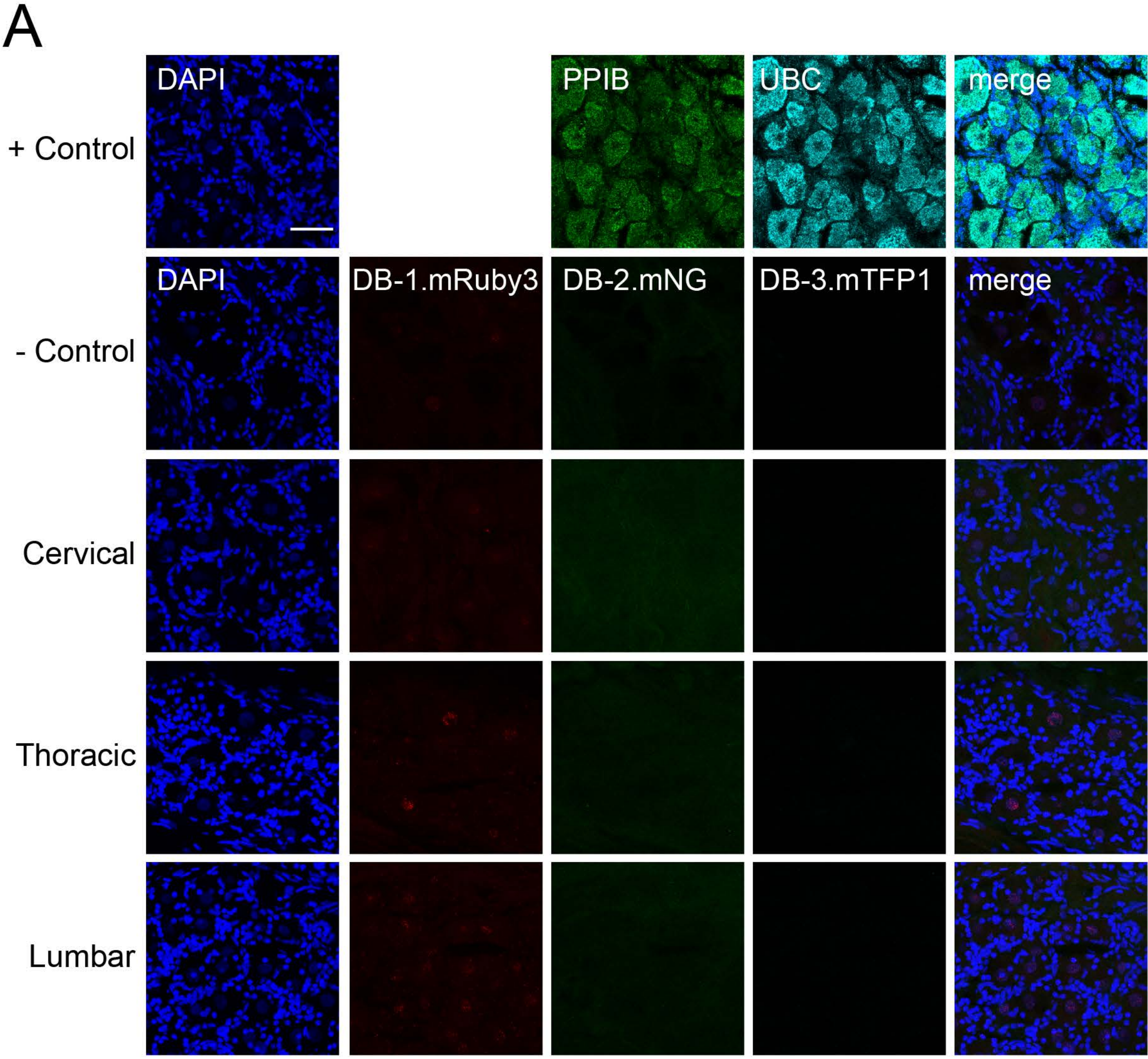


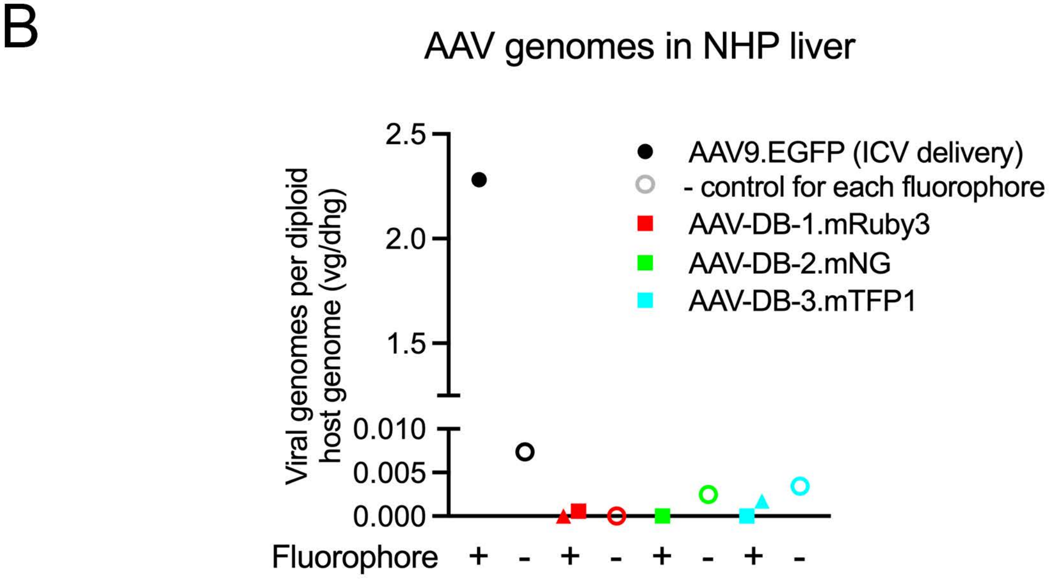

Supplemental Figure 1. Assessment of AAV-DB-1, AAV-DB-2 and AAV-DB-3 expression in off-targets tissues. A) FISH analyses for mRuby3, mNG and mTFP1 transcripts at cervical, thoracic, and lumbar NHP DRGs 24 days after AAV-DB intraparenchymal injection. Housekeeping transcripts PPIB and UBC were used as positive controls for FISH. B) Quantification of AAV genomes in liver of NHPs that received AAV-DB capsids by GP injection. Liver from a NHP injected ICV with AAV9-eGFP is included as a positive control.

Supplemental Figure 2

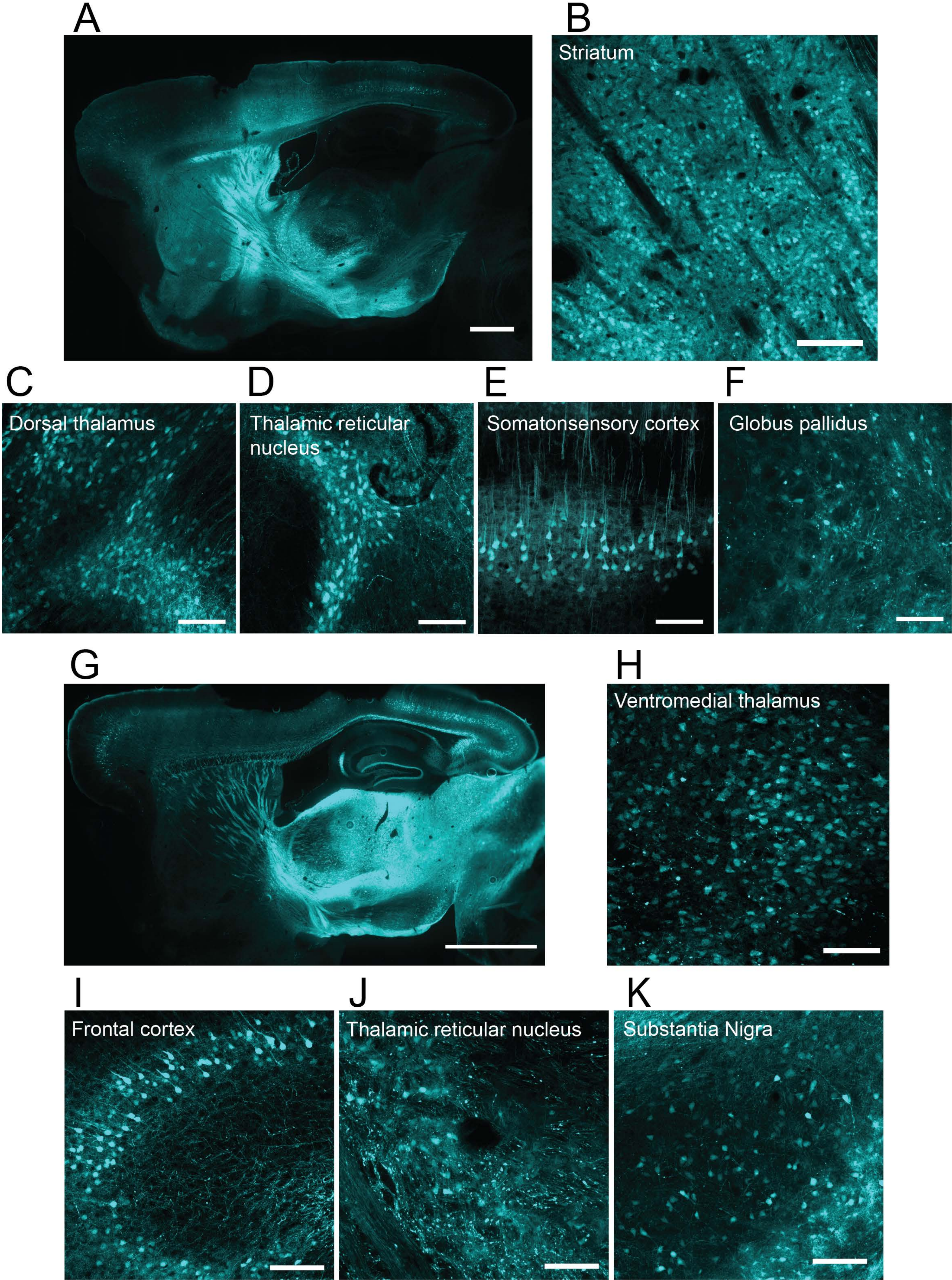

Supplemental Figure 2. Distribution of AAV-DB-3 following direct striatal or thalamic injection. A) Whole sagittal sections showing broad neuronal distribution of AAV-DB-3 after striatal delivery. B) mTFP1-positive neurons in MSNs in striatum. Retrograde positive neurons in the dorsal (C) and reticular (D) thalamus. Transgene positive neurons in the somatosensory cortex (E) and GPe (F). G) Whole section showing broad neuronal distribution of AAV-DB-3 after thalamic delivery. Positive labeled neurons located in the ventromedial (H) and reticular nucleus of the thalamus (J). I) mTFP1 positive neurons in the frontal association cortex. K) Retrograde labeled substantia nigra neurons. Scale bars: A) 1.5 mm, B-E) 200 µm.
